# Model Selection in Occupancy Models: Inference versus Prediction

**DOI:** 10.1101/2022.03.01.482466

**Authors:** Peter S. Stewart, Philip A. Stephens, Russell A. Hill, Mark J. Whittingham, Wayne Dawson

## Abstract

Occupancy models are a vital tool for applied ecologists studying the patterns and drivers of species occurrence, but their use requires a method for selecting between models with different sets of occupancy and detection covariates. The information-theoretic approach, which employs information criteria such as Akaike’s Information Criterion (AIC) is arguably the most popular approach for model selection in ecology and is often used for selecting occupancy models. However, the information-theoretic approach risks selecting models which produce inaccurate parameter estimates, due to a phenomenon called collider bias. Using simulations, we investigated the consequences of collider bias (using an illustrative example called M-bias) in the occupancy and detection processes of an occupancy model, and explored the implications for model selection using AIC and a common alternative, the Schwarz Criterion (or Bayesian Information Criterion, BIC). We found that when M-bias was present in the occupancy process, AIC and BIC selected models which inaccurately estimated the effect of the focal occupancy covariate, while simultaneously producing more accurate predictions of the site-level occupancy probability. In contrast, M-bias in the detection process did not impact the focal estimate; all models made accurate inferences, while the site-level predictions of the AIC/BIC-best model were slightly more accurate. Our results demonstrate that information criteria can be used to select occupancy covariates if the sole purpose of the model is prediction, but must be treated with more caution if the purpose is to understand how environmental variables affect occupancy. By contrast, detection covariates can usually be selected using information criteria regardless of the model’s purpose. These findings illustrate the importance of distinguishing between the tasks of parameter inference and prediction in ecological modelling. Furthermore, our results underline concerns about the use of information criteria to compare different biological hypotheses in observational studies.

**Open Research Statement:** Code to fully reproduce our simulations and analyses is available at: https://zenodo.org/badge/latestdoi/462801230

## Introduction

The patterns and drivers of species occurrence are of fundamental interest to ecologists. Predicting where species occur enables ecologists to tackle key problems such as understanding the spread of invasive species (Gormley *et al*. 2011), assessing the efficacy of protected areas (Midlane *et al*. 2014), and evaluating the extinction risk (Breiner & Bergamini 2018) and recovery of populations and species (Akçakaya *et al*. 2018). Understanding the drivers of occurrence is also important; interventions to mitigate the factors which threaten species must be informed by the diagnosis of those factors (Caughley 1994). Many studies have aimed to determine how occurrence is driven by factors including forest degradation (Zimbres *et al*. 2018), agricultural expansion (Semper-Pascual *et al*. 2020), wildfires (Hossack *et al*. 2013), and anthropogenic noise pollution (Chen & Koprowski 2015).

A key challenge in studying the patterns and drivers of occurrence is that ecologists are often constrained to the use of observational data; experimental manipulations of ecological systems may be physically impossible, logistically unfeasible, or unethical. Consequently, one approach is to use a model which relates observed variation in species occurrence to one or more environmental covariates. The model can then be used to predict species occurrence at new sites, or to examine the effect of each covariate on occurrence (Shmueli 2010). Occupancy models are often used for this purpose because they deal with imperfect detection (MacKenzie *et al*. 2002, 2006). They do so by modelling the probability that a species occupying the site is detected, often including environmental covariates to explain variation in detectability among sites (MacKenzie *et al*. 2002, 2006). Occupancy models therefore contain one set of covariates for occupancy probability, and a second set for detection probability; the challenge is to select suitable sets of covariates to include in the model. This challenge can be framed as a problem of model selection (Robins & Greenland 1986; Buckland *et al*. 1997; Forster 2000; Burnham & Anderson 2004; Johnson & Omland 2004).

One approach to model selection is the information-theoretic approach (Anderson *et al*. 2000; Burnham & Anderson 2001, 2004; Lukacs *et al*. 2007; Burnham *et al*. 2011), which compares models in terms of their relative Kullback-Leibler (KL) divergence – the relative distance between each model and “full reality”, in units of information entropy (Forster 2000; Burnham & Anderson 2001; McElreath 2021). Information criteria, of which Akaike’s information criterion (AIC; Akaike 1973) is the most commonly used, estimate the relative KL divergence of each model using the sample data (McElreath 2021). AIC is calculated for each model by taking the in-sample deviance (a measure of how well the model fits the data), and adding an overfitting penalty term proportional to the number of parameters in the model (Akaike 1973; Burnham *et al*. 2011). Consequently, AIC favours parsimonious models which balance underfitting and overfitting, with the aim of producing better out-of-sample predictions (McElreath 2021).

Some proponents of the information-theoretic approach argue that each model in the candidate set should represent a different biological hypothesis, and that the models’ relative AIC scores indicate the strength of evidence for each hypothesis (Burnham *et al*. 2011). However, insights from the field of causal inference reveal a potential problem with this approach – collider bias. Collider bias is a form of confounding, where the estimate of an effect is biased due to some feature of the data-generating process (Greenland 2003; Greenland & Mansournia 2015). However, unlike the classical notion of confounding, collider bias arises not from omitting an important variable from the statistical model (omitted variable bias; Clarke 2005), but from including a variable that leads to bias (sometimes referred to as a “bad control”; Cinelli *et al*. 2020; McElreath 2021). In the simplest case, collider bias arises from including a variable which is a function of both the explanatory and response variable of interest (Greenland 2003). For example, in spotted hyenas (*Crocuta crocuta*), reproductive state is likely a function of both social connectedness and immune function: therefore, including reproductive state as a covariate will induce a spurious association between social connectedness and immune function (Laubach *et al*. 2021). In more complex examples with more than three variables, collider bias can arise from the relationships between variables without the response variable directly affecting any other variable in the system (Fig. 1).

**Figure 1.**
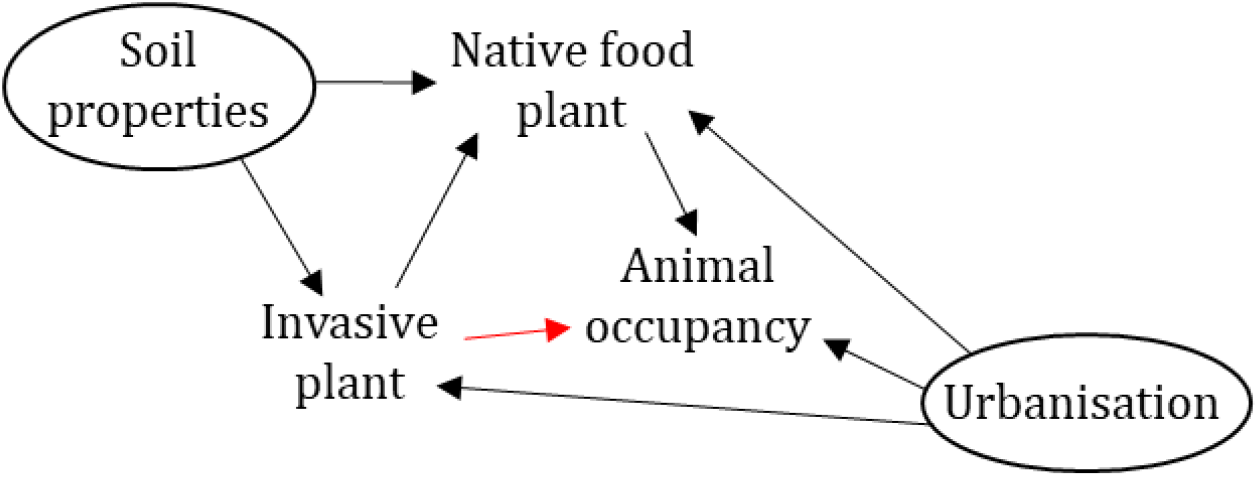
Directed acyclic graph showing a hypothetical example in which we are interested in estimating the direct effect of the density of an invasive plant on animal occupancy (red arrow). Invasive plant density also influences the density of a native food plant, which in turn influences the animal’s occupancy. Soil properties influence the density of both the native and invasive plant – these properties are latent (unobserved), and so are shown in a circle. Urbanisation influences the densities of both plant species, as well as animal occupancy – here, urbanisation is also latent. In this example, native food plant density is a collider which sits on the back-door path from invasive plant density to animal occupancy, *Invasive plant* ← *(Soil properties)* → *Native food plant* ← *(Urbanisation)* → *Animal occupancy*. Including native food plant density in the model opens the back-door path, biasing the estimated direct effect of the invasive plant. Notably, in this example the effect of interest cannot be estimated by including any of the available covariates – measuring and including urbanisation is necessary.

As AIC and other information criteria select models based on their expected predictive performance, they will not necessarily select models which return accurate parameter estimates (McElreath 2021). For example, Luque-Fernandez *et al*. (2019) demonstrated that for a simple collider example, including the collider covariate improved the model’s AIC score, while simultaneously resulting in an estimated effect size which was far from the true value. It is straightforward to extend their simulation to show that this result is consistent across a broad range of effect sizes (Fig. 2). However, the implications for model selection in occupancy modelling, in which covariates must be selected for both the occupancy and detection sub-models, have yet to be investigated.

**Figure 2.**
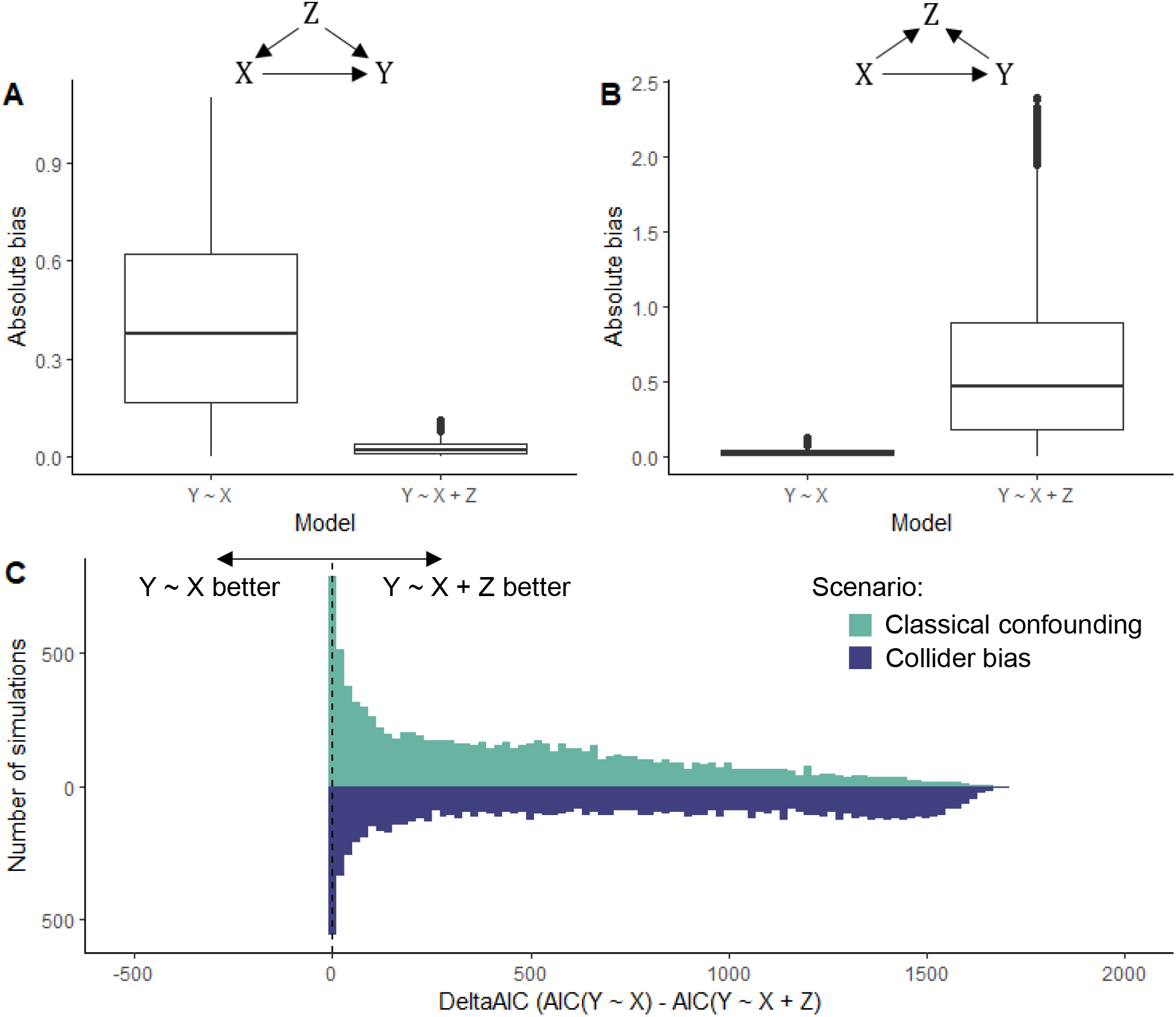
Luque-Fernandez *et al*. (2019) presented simulations illustrating classical confounding and collider bias in a simple linear model. We extended their example by conducting 10000 simulation iterations for each example, with each iteration using true effect sizes drawn from a uniform distribution between -2 and 2 to ensure that the results are not specific to the choice of parameter values. In the classical confounding example **(A)**, including the variable Z reduces the absolute bias (the absolute difference between the estimated and true effect: 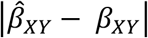 when estimating the effect of X on Y. Conversely, in the collider example **(B)**, including Z increases the absolute bias. However, in both cases Akaike’s Information Criterion (AIC) favours the model which includes Z **(C)**, illustrating that AIC does not always favour models which produce accurate parameter estimates. R code to reproduce the simulations is available at: https://zenodo.org/badge/latestdoi/462801230

To address this topic, we investigated the consequences of a form of collider bias known as “M-bias” (Fig. 3 ; Greenland 2003; Cinelli *et al*. 2020) in an occupancy modelling framework, and explored the implications for model selection using the information-theoretic approach (using AIC). We also examined the performance of a common alternative to AIC, the Schwarz criterion (or Bayesian Information Criterion, BIC; Schwarz 1978), which some authors suggest can be used for selecting the “true” model from the candidate set (Aho *et al*. 2014). We focused on M-bias because it is a common illustrative example in the causal inference literature (*e*.*g*. Greenland 2003; Cinelli *et al*. 2020), which demonstrates several important points: i) conditioning on a variable may cause more harm than omitting it from the model (Greenland *et al*. 1999), ii) collider bias can occur even when none of the variables considered are directly affected by the response variable, and iii) the importance of considering potential latent (unobserved) variables when drawing inferences. In our simulation-based approach, we generated datasets where M-bias was present in the occupancy process, the detection process, or both. We then fitted occupancy models with different sets of covariates to these datasets, and evaluated them on the accuracy of their parameter inferences, the accuracy of their site-level occupancy predictions, and their level of support from AIC and BIC.

**Figure 3.**
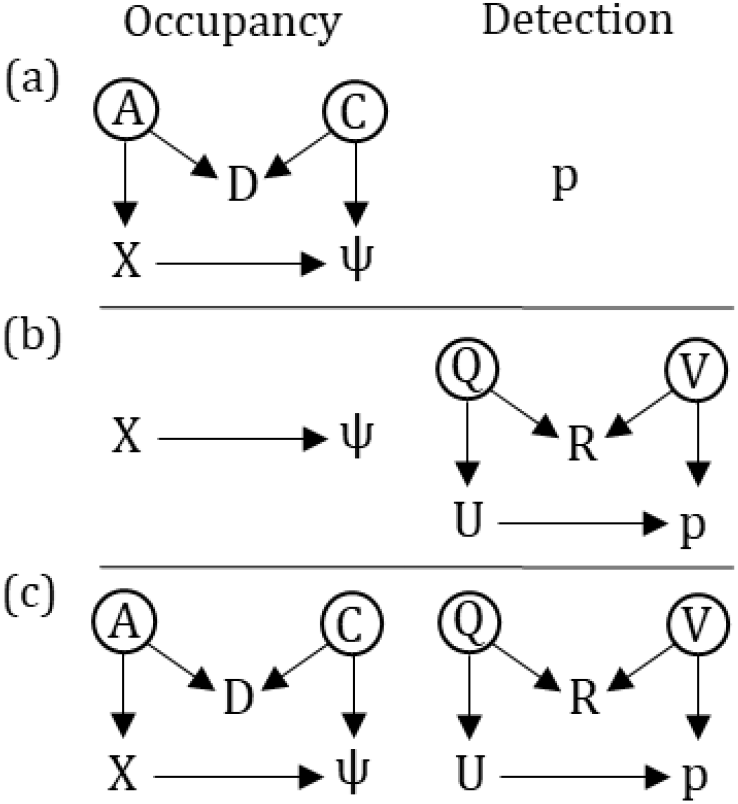
Directed acyclic graphs (DAGs) showing the relationship between variables in the occupancy and detection processes of the three simulation scenarios: **a)** scenario 1 (M-bias in the occupancy process), **b)** scenario 2 (M-bias in the detection process), **c)** scenario 3 (M-bias in both the occupancy and detection processes). *ψ* represents the probability that a site is occupied by a species (occupancy probability), while *p* represents the probability that a species is detected given that it is present at a site (detection probability). *X* represents an explanatory variable for which we are interested in estimating the effect on the occupancy probability. *A, C, D, Q, U, R* and *V* represent additional variables which are thought to be important in the system. Circled variables are latent (unobserved).

## Methods

### Causal inference, directed acyclic graphs and M-bias: a primer

Causal inference is concerned with predicting the consequences of intervening in a system, as well as inferring counterfactual outcomes – events which might have happened, under hypothetical unrealised conditions (Pearl 2009; Pearl *et al*. 2016). Importantly, causal inference is not about ‘inferring causation from correlation’ – conclusions about causality cannot be made from the data alone, but require causal assumptions about the process which generated the data (Pearl 2009; Pearl *et al*. 2016). These assumptions, rather than remaining implicit, should be communicated clearly so that they are open to scrutiny, debate, sensitivity analysis, and verification (Pearl 2009).

Causal assumptions are typically expressed in a graphical model, usually a directed acyclic graph (DAG) where the nodes represent variables and the edges represent the relationships between them (Pearl 1995). The nodes on a DAG can represent both observed and latent variables – the latter are usually shown surrounded by a circle (*e*.*g*., A and C in Fig. 3a), and we show them surrounded by brackets in the main text. The edges linking each node are directional, and are thus represented as arrows; these arrows represent the mechanistic links between the variables (Greenland *et al*. 1999). The succession of edges linking one variable to another, regardless of which direction these edges are pointing in, is known as a path (Pearl 1995). For instance, Fig. 3a has two paths connecting *X* and *ψ*: *X* → *ψ*, and *X* ← *(A)* → *D* ← *(C)* → *ψ*.

A DAG corresponds to a non-parametric structural equations model (Pearl 1995), where each node is represented as a function of the variables which have edges going into it, as well as a “disturbance term” (ε) which represents the effect of omitted variables which are assumed to be independent of one another (Pearl 1995). For example, the DAGs in Fig. 3c correspond to the following set of equations:

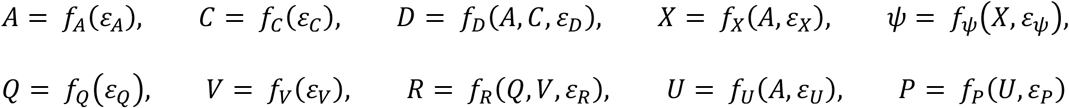

Importantly, the model is non-parametric in the sense that the form of these functions does not have to be specified (Pearl 1995; Greenland *et al*. 1999), allowing DAGs to accommodate non-linear functions and interactions. For example, the variable *D* in the above equation could be modelled as a nonlinear function of A and C, or a linear interaction between A and C.

Once a DAG has been specified, it is analysed to identify the set of variables necessary to remove structural confounds from the effect of interest. One strategy is to condition on (*i*.*e*., include as covariates) the variables which satisfy the “back-door criterion”, in which the aim is to “close all back-door paths” linking the explanatory and response variable of interest (Pearl 1995). A back-door path is defined as any path which has an arrow entering the focal explanatory variable (Pearl 1995) – for example, in Fig. 3a the path *X* ← *(A)* → *D* ← *(C)* → *ψ* is a back-door path.

By default, paths which are a “fork” (*e*.*g*., X ← Z → Y) or “pipe” (*e*.*g*., X → Z → Y) are open, and conditioning on the middle variable (Z) closes them (Greenland 2003; Pearl *et al*. 2016; McElreath 2021). Paths which are a collider (*e*.*g*., X → Z ← Y) are closed by default, and conditioning on the middle variable (Z) opens the path (Greenland *et al*. 1999; Greenland 2003; Pearl *et al*. 2016; McElreath 2021). A back-door path only needs to be closed in one place; for instance, X ← (A) → D ← (C) → ψ is closed because it is blocked by the collider at D, irrespective of the forks at A and C.

The path structure of a given DAG implies that specific pairs of variables will be independent of one another, conditional on a set of other variables which close the paths between them. Examining whether these conditional independences hold in a given dataset is an important aspect of analysing whether a DAG is consistent with the observed data (Textor *et al*. 2016).

The DAG displayed for *ψ* in Fig. 3a, for example, contains an “M”-shaped structure, which can induce a type of confounding known as M-bias (Greenland 2003). By conditioning on the collider (D in Fig. 3a), the back-door path is opened and the effect of interest (X on ψ in Fig. 3a) is confounded. When the back-door path contains latent variables (A and C in Fig. 3a) it is not possible to condition on them to close the path again because they are unobserved by definition. If the objective is to obtain an unconfounded estimate of the effect of X on ψ, then the correct approach is to avoid conditioning on D.

### Simulation study

To explore the effects of M-bias in both the occupancy and detection components of an occupancy model, we conducted simulations of three different scenarios (Fig. 3) in which the target for inference was the effect of variable *X* on occupancy probability (*ψ*). In the first scenario (Fig. 3a), *ψ* was part of an M-graph while the detection probability (*p*) was fixed at 0.5. In the second scenario (Fig. 3b), *ψ* depended only on *X*, and *p* was now part of an M-graph. In the final scenario (Fig. 3c), both the occupancy and detection probabilities were part of M-graphs.

All three simulations followed the same process: **1)** generate a dataset with known parameter values, using the relationships between variables embodied in the relevant DAGs (Fig. 3); **2)** fit a number of occupancy models to the dataset (Table 1); **3)** evaluate each model’s accuracy in parameter estimation and prediction; **4)** evaluate each model’s quality under the information-theoretic framework. Each simulation was repeated 1000 times. We conducted our simulations in R (v 4.0.5; R Core Team, 2021), and provide code to reproduce our simulations and analyses at: https://zenodo.org/badge/latestdoi/462801230

**Table 1.** Occupancy models fitted in each simulation scenario. All models also included an intercept term for both occupancy and detection. *X* represents an explanatory variable for which we are interested in estimating the effect on the occupancy probability. *D, U*, and *R* represent additional variables which are thought to be important in the occupancy or detection processes.

| **Model** | **Occupancy covariates** | **Detection covariates** |
| --- | --- | --- |
| *Scenario 1* |  |  |
| Model 1 | $X$ | n/a |
| Model 2 | $X + D$ | n/a |
| *Scenario 2* |  |  |
| Model 1 | $X$ | $U$ |
| Model 2 | $X$ | $U + R$ |
| *Scenario 3* |  |  |
| Model 1 | $X$ | $U$ |
| Model 2 | $X + D$ | $U$ |
| Model 3 | $X$ | $U + R$ |
| Model 4 | $X + D$ | $U + R$ |

#### 1) Generating a dataset

Data were simulated for 3000 sites with 40 surveys each. These values were deliberately high to ensure that any inaccuracy was not primarily driven by an underpowered design.

We first drew effect sizes for each arrow in the relevant DAG from a uniform distribution between -1 and 1. Values for explanatory variables with no ingoing arrows on the DAG were then drawn from a normal distribution with a mean of 0 and standard deviation of 1. We then generated values for the other explanatory variables from the appropriate variables and effect sizes (*i*.*e*., those from ingoing arrows on the DAG; Fig. 3), plus a “disturbance term” (*sensu* Pearl 1995) drawn from a normal distribution with a mean of zero and standard deviation of 0.025. The disturbance term was required to be non-zero to avoid perfect collinearity between the explanatory variables which would result in model failure.

We then generated the true occupancy and detection probabilities. The log-odds of occupancy and detection were first calculated as a linear function of the explanatory variables with ingoing arrows on the DAG (Fig. 3), and the inverse-logit was then taken to obtain the probability. The true occupancy state of each site was then simulated as a Bernoulli trial with probability of success equal to the occupancy probability. We assumed that the true occupancy state remained constant for each site, which is a key assumption of single-season occupancy models (MacKenzie *et al*. 2002).

Detection histories were then generated for each site, as 40 Bernoulli trials with probability of success equal to the true occupancy state multiplied by the detection probability. This means that the species was never observed at sites in which it was absent (occupancy state = 0), while when the species was present (occupancy state = 1) it had a chance of being observed on each visit, with probability equal to the detection probability. The detection histories for each site represent the observed data that would be collected by an ecologist in the field, alongside the covariate values for the non-latent variables.

#### 2) Fitting models

Occupancy models were fitted to each dataset using the *occu* function in the R package *unmarked* (v.1.0.0; Fiske & Chandler, 2011), which implements the single-season occupancy model developed by MacKenzie and colleagues (2002). The models used the logit link function because we used the inverse-logit to simulate the data. We fitted models with various combinations of observed variables (*i*.*e*., excluding latent variables) for each scenario (Table 1). All models included the focal covariate for occupancy (*X*). In scenario 1, detection probability was modelled as an intercept only, as it did not depend on any covariates.

#### 3) Evaluating Model Performance

In each scenario, all models were evaluated on the accuracy of their parameter inferences and predictions. To quantify how accurately each model estimated the effect of covariate *X* on the occupancy probability *ψ*, we calculated the bias and absolute bias:

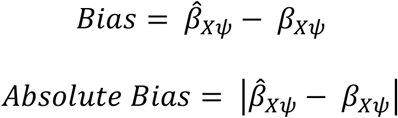

Where 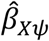 and *β*_*Xψ*_ are the estimated and true effects of *X* on *ψ*, respectively. Additionally, we constructed 95% confidence intervals for 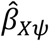 as follows:

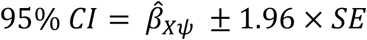

where SE is the standard error of the 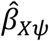 estimate, obtained from the model summary table. We then checked whether the true value, *β*_*Xψ*_, was found within this interval. We also checked whether the sign (positive or negative) of 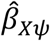 was the same as that of *β*_*Xψ*_.

To evaluate each model’s predictions, we used the *predict* function in R to make predictions for the occupancy probability value for each site in two datasets. We first made predictions for the data to which the model was fitted, to examine how the model retrodicted the sample. We then examined the model’s performance in out-of-sample prediction by making predictions for a new dataset (also 3000 sites with 40 surveys each), which was generated using the same true parameter values as the original dataset. This corresponds to the common task in which occupancy models are fitted to one set of data (*e*.*g*., grid squares sampled from a region) and used to extrapolate to a wider set of data which are assumed to arise from the same data-generating process (*e*.*g*., other grid squares in the region).

To assess the accuracy of the model retrodictions and predictions, we first calculated the error for each site:

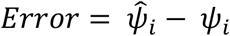

where 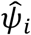 and *ψ*_*i*_ are the estimated and true occupancy probabilities for site *i*, respectively. To summarise this error over all sites in the dataset, we calculated the mean absolute error:

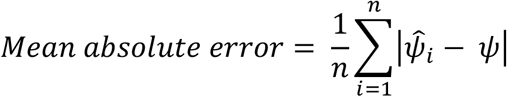

where *n* is the number of sites. Additionally, we checked whether the true occupancy probability for each site was within the 95% confidence interval obtained from the prediction, and calculated the proportion of sites for which this was the case.

#### 4) Evaluating models under the information-theoretic framework

To examine the degree of support for each model under the information-theoretic framework we obtained the AIC value for each model *m* from the model’s summary table. We then calculated the ΔAIC value for each model in the scenario as:

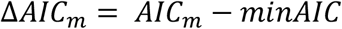

where *AIC*_*m*_ is the AIC value for model *m*, and *minAIC* is the lowest AIC value for the set of models in the scenario.

Proponents of the information-theoretic approach have advocated for multimodel inference (*e*.*g*., Anderson *et al*. 2000; Burnham & Anderson 2004; Burnham *et al*. 2011), in which inferences are made using the entire candidate set of models, each weighted using Akaike weights derived from AIC. This approach accounts for uncertainty in the model selection process (Burnham & Anderson 2004). We calculated Akaike weights (*w*) for each model *m* as:

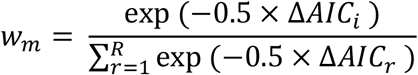

where *R* is the number of models in each scenario.

We also considered the Schwarz criterion (BIC; Schwarz 1978) as an alternative to AIC. BIC is defined as:

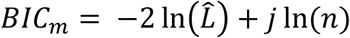

where 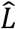 is the maximum likelihood estimate and thus −2 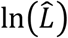 the in-sample deviance, *j* is the number of parameters, and *n* is the number of observations in the dataset. We calculated BIC, ΔBIC, and BIC weight values for each model using the R package *AICcmodavg* (Mazerolle, 2020). ΔBIC and BIC weights are defined as for ΔAIC and AIC weights, but using the BIC values for each model.

## Results

### Scenario 1: M-Bias in the Occupancy Process

When M-bias was present in the occupancy process, model 1 (*ψ* ∼ *X*) estimated the true effect of *X* on *ψ* much more accurately than model 2 (*ψ* ∼ *X + D*) (Table 2, Fig. 4a,b). However, comparing the two models’ predictive accuracy showed the opposite picture; model 1 generally produced worse predictions than model 2 (Table 2, Fig. 5a,b). The models’ retrodictive accuracy was also qualitatively similar to their predictive accuracy (Fig. S1). AIC and BIC both showed clear support for model 2 in the majority of simulations (Fig. 6a,b); in 85.1% of simulations model 2 received an Akaike weight of >0.99, and in 63.0% of simulations it received the entire weight (Fig. 6b). The few simulations in which model 1 received more Akaike weight were mostly those in which β_Cψ_ was small (Fig. S2). A similar pattern of results was observed for BIC (Fig. S2), although when BIC assigned weight to model 1 it generally assigned more weight than AIC (Fig. 6a).

**Table 2.**
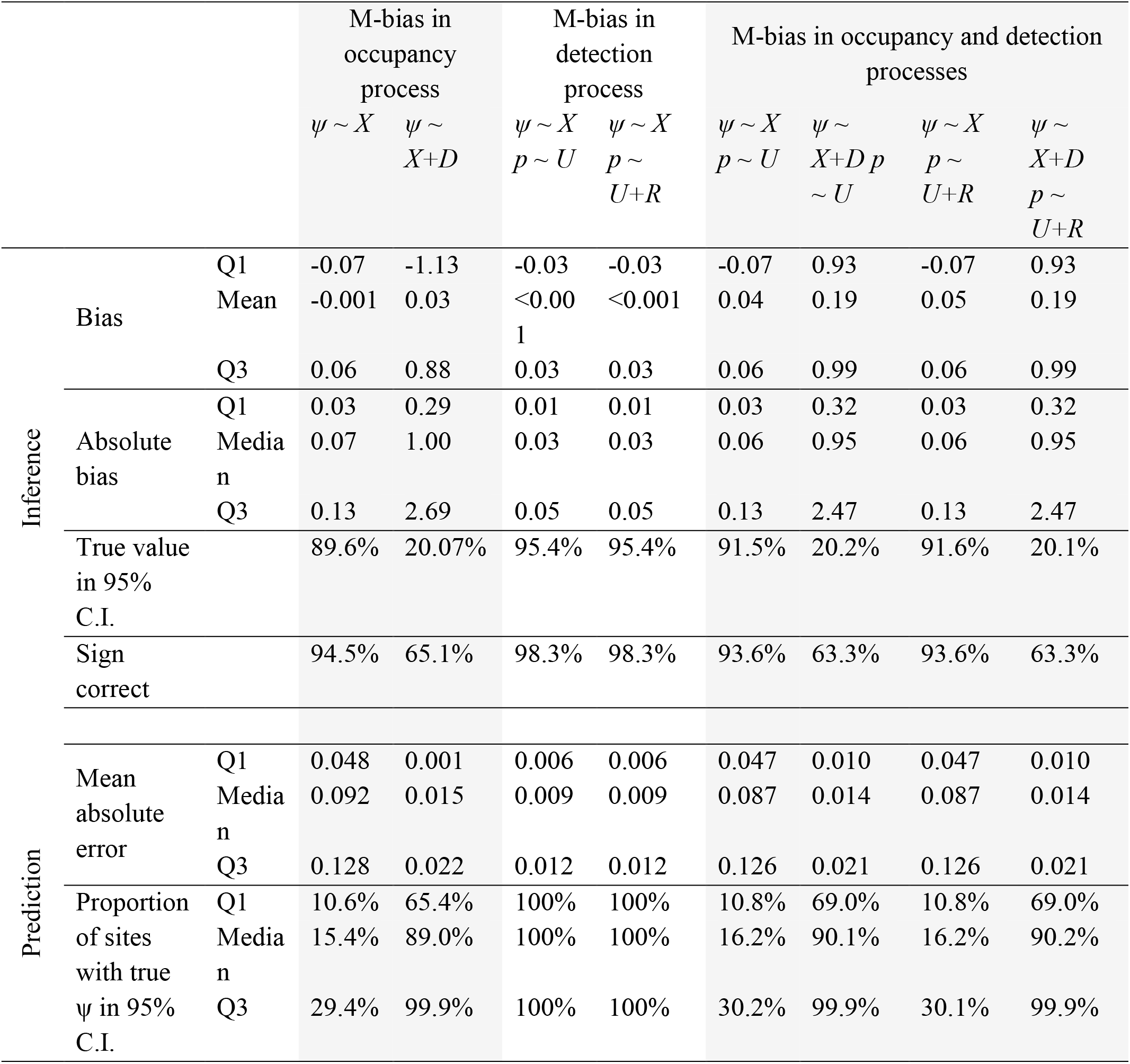
Inferential and predictive accuracy of occupancy models across all simulations (n=1000) for each of the three scenarios: M-bias in the occupancy process only (left), M-bias in the detection process only (centre), and M-bias in both the occupancy and detection processes (right). Each model is shown as a separate column, with the occupancy *(ψ)* and detection *(p)* covariates displayed at the top. Each row indicates a separate metric summarising the models’ performance across the simulations. Full definitions of each metric can be found in the main text.

**Figure 4.**
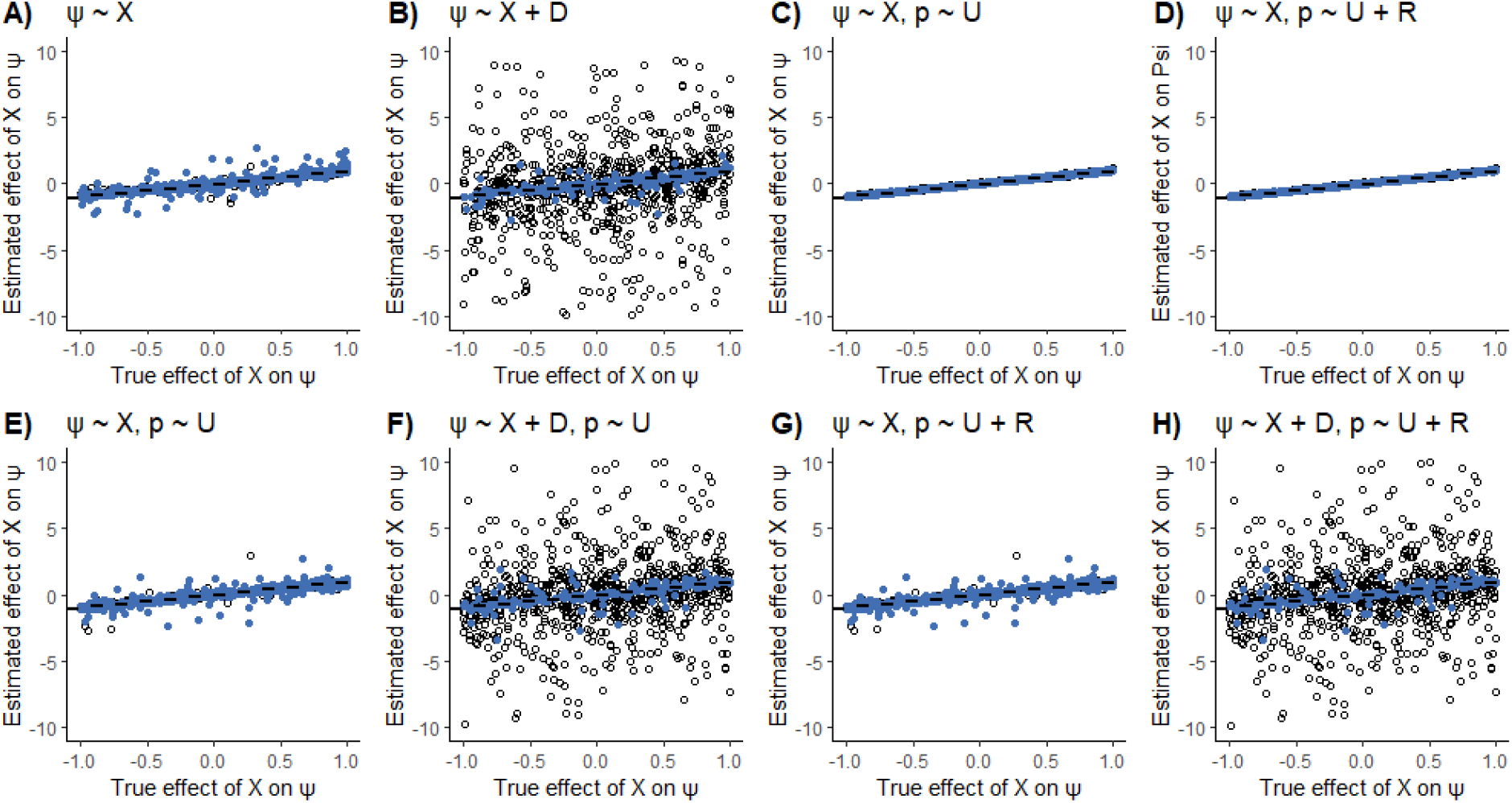
True versus estimated effect of *X* on occupancy probability (*ψ*), for the following occupancy models: **A)** scenario 1, model 1; **B)** scenario 1, model 2; **C)** scenario 2, model 1; **D)** scenario 2, model 2; **E)** scenario 3, model 1; **F)** scenario 3, model 2; **G)** scenario 3, model 3; **H)** scenario 4, model 4. Each point represents the result from one simulation, with 1000 simulations in total. Note that the y-axis is truncated at -10 and 10; plots B, F, and H omit 40, 38 and 38 points respectively which lay outside this range. Blue points indicate that the true value was contained within the estimate’s 95% confidence interval, while unfilled circles indicate that the true value was not contained within the interval. Dashed black line indicates equality between the true and estimated effect – increased vertical distance from this line indicates a more biased estimate. Each model’s covariates for *ψ* and the detection probability (*p*) are shown above their respective plot. For explanation of the scenarios, see Table 1 and main text.

**Figure 5.**
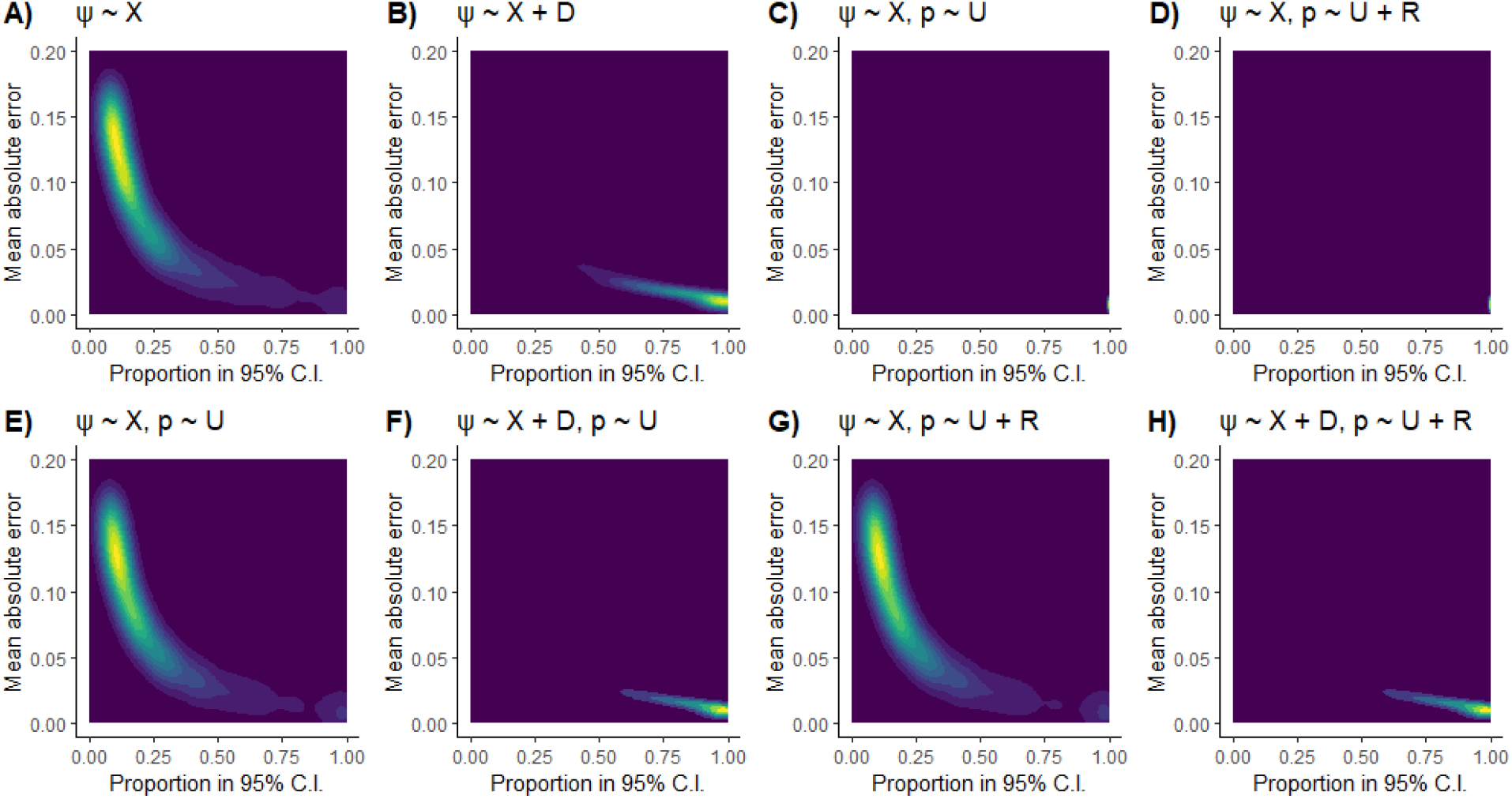
Kernel density estimate contours showing two measures of predictive accuracy when predicting site-level occupancy probability (*ψ*), for 1000 simulations. The x-axis shows the proportion of sites (out of 3000 sites) for which the true occupancy probability was contained within the 95% confidence interval around the model’s prediction. The y-axis shows the mean absolute error of the predictions. Consequently, the bottom right of each plot indicates higher predictive accuracy, while the top left indicates lower predictive accuracy. The density of simulations within this area is shown by the coloured contours, with lighter colours indicating a higher density of simulations. Results are displayed for the following occupancy models: **A)** scenario 1, model 1; **B)** scenario 1, model 2; **C)** scenario 2, model 1; **D)** scenario 2, model 2; **E)** scenario 3, model 1; **F)** scenario 3, model 2; **G)** scenario 3, model 3; **H)** scenario 4, model 4. Each model’s covariates for *ψ* and the detection probability (*p*) are shown above their respective plot. For explanation of the scenarios, see Table 1 and main text.

**Figure 6.**
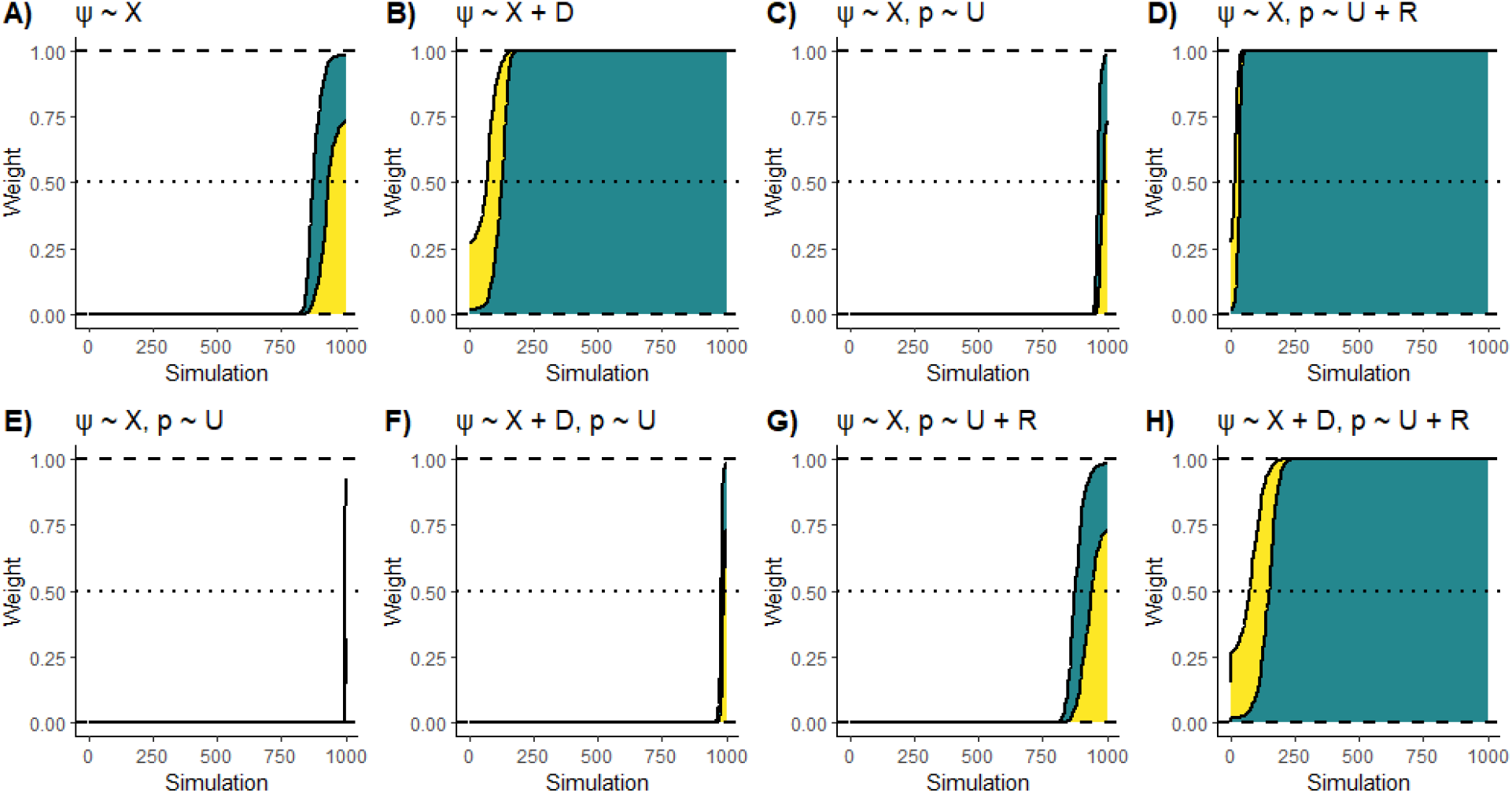
Akaike weight (yellow area) and BIC weight (blue area) for 1000 simulations of eight occupancy models. Simulations are shown ranked by weight, with higher Akaike and BIC weights shown on the right. Higher weights correspond to lower AIC/BIC values, indicating more support for the model. The panels display: **A)** scenario 1, model 1; **B)** scenario 1, model 2; **C)** scenario 2, model 1; **D)** scenario 2, model 2; **E)** scenario 3, model 1; **F)** scenario 3, model 2; **G)** scenario 3, model 3; **H)** scenario 4, model 4. Each model’s covariates for the occupancy probability (*ψ*) and the detection probability (*p*) are shown above their respective plot. For explanation of the scenarios, see Table 1 and main text. Dashed horizontal lines are shown for weights of 0, 0.5, and 1.

### Scenario 2: M-Bias in the Detection Process

When M-Bias was present in the detection process, both models 1 (*ψ* ∼ *X, p* ∼ *U*) and 2 (*ψ* ∼ *X, p* ∼ *U + R*) accurately estimated the effect of *X* on *ψ* (Table 2, Fig. 4c,d). Both models also made very accurate predictions (Table 2, Fig. 5c,d), although model 2 performed slightly better - in 86.4% of simulations, the true occupancy probability was contained within model 2’s 95% confidence interval for every predicted site, whereas model 1 accomplished this in 85.8% of simulations. Similar results were observed for the models’ retrodictive accuracy (Fig. S1c,d). Both AIC and BIC assigned the majority of weight to model 2 in most simulations (Fig. 6c,d).

### Scenario 3: M-Bias in the Occupancy and Detection Processes

When M-Bias was present in both the occupancy and detection processes, models 1 (*ψ* ∼ *X, p* ∼ *U*) and 3 (*ψ* ∼ *X, P* ∼ *U + R*) estimated the effect of X on ψ much more accurately than models 2 (*ψ* ∼ *X + D, P* ∼ *U*) and 4 (*ψ* ∼ *X + D, P* ∼ *U + R*) (Table 2, Fig. 4e-h). In general, the 95% confidence interval around the estimate in models 2 and 4 only contained the true value when *β*_*AD*_ and *β*_*Cψ*_ (and to a lesser extent *β*_*AX*_) were relatively small (Figs. S4,5). In contrast, models 2 and 4 made more accurate predictions than models 1 and 3 (Table 2, Fig. 5e-h), and similar results were obtained for retrodictive accuracy (Fig. S1e-h). Both AIC and BIC showed clear support for model 4 in the majority of simulations (Fig. 6h); the model received an Akaike weight of >0.99 in 82.2% of the simulations. While model 3 did occasionally receive weight, this mostly occurred when β_Cψ_ was small (Fig. S6) and the model still never received the entire weight (Fig. 6g). BIC weights were similar to the Akaike weights, although BIC assigned more weight to model 3 in some simulations (Fig.4g). Again, these instances were generally when *β*_*Cψ*_ was small (Fig. S6).

## Discussion

Here, we investigated the consequences of M-bias (a specific form of collider bias) for occupancy modelling, and explored the implications for model selection using AIC and BIC. In our simulations, we observed that when M-bias was present in the occupancy process, AIC and BIC favoured a model which produced a highly inaccurate estimate of the focal effect but produced more accurate predictions and retrodictions of the site-level occupancy probability. In contrast, M-bias in the detection process did not result in inaccurate estimates of the focal effect – both models made accurate inferences. Both models also made accurate predictions and retrodictions, although those of the AIC/BIC-best models were slightly better. The same results were observed when M-bias was present in both the occupancy and detection processes: the model favoured by AIC and BIC produced inaccurate inferences but more accurate predictions, while models made similarly accurate inferences regardless of M-bias in the detection process. These results have important implications for model selection in occupancy models, as well as for how the information-theoretic approach is applied in ecological modelling more generally.

### Information criteria select models which produce poor parameter inferences, but good predictions

When M-bias was present in the occupancy process, the model which received the greatest support from AIC and BIC produced highly inaccurate estimates of the effect of the variable of interest *(X)* on the occupancy probability *(ψ)*. Such biased estimates are not informative about the biological drivers underlying the observed pattern, nor do they accurately predict the consequences of intervening in the system. To illustrate the importance of this result, imagine that the fourth iteration of the first scenario, in which M-bias is present in the occupancy process only, represents a real ecological study where the objective is to inform management to increase the occupancy of a species. In this example, model 2 (*ψ* ∼ *X + D*) received Akaike and BIC weights of one – it is clearly the “best” model, according to these criteria. The model estimated the effect of *X* on *ψ* as -1.22 (± 0.12 standard error) – a strong negative effect on occupancy. However, this effect is confounded; in reality, the effect of *X* on *ψ* is strongly positive (0.72). Furthermore, the estimated effect of *D* on *ψ* is also strongly negative (−1.70 ± 0.09), when *D* actually has zero effect on occupancy. In this example, the AIC/BIC “best” model suggests that to increase *ψ*, we should intervene to reduce *X* and *D*. In fact, this would reduce occupancy, as well as waste resources managing the unrelated variable *D*. If applied to a real conservation problem informed by occupancy models (*e*.*g*. Hossack *et al*. 2013; Chen & Koprowski 2015; Zimbres *et al*. 2018; Semper-Pascual *et al*. 2020), the results could be disastrous. Such poor inferences were the norm in our simulations; confounded models were only able to estimate the direction of the focal effect correctly in 65.1% of cases at best – little better than the accuracy we would expect from guessing.

While the models supported by AIC and BIC produced biased parameter estimates, they also produced more accurate predictions and retrodictions of the occupancy probability at each site. This is because these models include the variable *D* which has an open path to *ψ*; including *D* provides additional information about the variation in *ψ*, improving prediction. However, because *D* is a collider, including it opens the back-door path from *X* to *ψ*, biasing the estimated effect of *X*. From the perspective of AIC and BIC, including *D* results in a reduced in-sample deviance which typically outweighs the penalty for adding the additional variable; this reduction must be greater to outweigh BIC’s larger penalty term, which is why BIC was more conservative in its tendency to select confounded models in our simulations (Fig. 6). This also explains why AIC and BIC tended to prefer the non-confounded model (omitting *D*) when *β*_*Cψ*_ was close to zero (Figs. S2, S6); the near-zero effect of *C* on *ψ* meant that the path from *D* to *ψ* through *C* was almost blocked (the other path from *D* to *ψ* was blocked by conditioning on *X*), and therefore *D* explained relatively little variation in *ψ*.

In contrast to the effects of M-bias in the occupancy process, M-bias in the detection process did not affect inferences about the effect of *X* on *ψ*. Additionally, including the collider variable *R* in the detection sub-model slightly improved the accuracy of the model’s predictions of the site-level occupancy probability. As for the occupancy process example, these results can be explained by considering how the path structure between variables will affect the change in deviance when a variable is included; as the variable *R* has an open path to *p*, including *R* explains additional variation in the detection probability, reducing the deviance and allowing the model to account better for imperfect detection when estimating the occupancy probability. As the detection probability is generally regarded as a nuisance parameter (Karavarsamis 2015), it is inconsequential that the effect of the other detection covariate (U) will be confounded. Therefore, as long as the aim is to explain as much variation in the detection probability as possible, information criteria can be used for selecting detection covariates.

The tendency for information criteria to favour confounded models with greater predictive ability is not confined to collider bias. For example, simulations by McElreath (2021) showed that information criteria tend to select models which condition on the mediator (*M*) in a pipe (*e*.*g*., *X* → *M* → *ψ*), inducing post-treatment bias (Rosenbaum 1984). Information criteria prefer models which include *M* because *M* has an open path to *ψ*, so including *M* explains additional variation in *ψ* (McElreath, 2021). However, including *M* also confounds the estimated effect of *X* – not by opening a non-causal back-door path as in the collider example, but instead by closing the causal front-door path which runs through *M* (McElreath, 2021). Therefore, if our aim is to estimate the effect of *X* on *ψ* then we should omit *M* from the model, whereas if our sole aim is to predict the value of *ψ* then including *M* would improve these predictions. We also expect these results to apply in other scenarios, such as case control bias (Cinelli *et al*. 2020).

### Inference and prediction are separate tasks

The key point supported by our results is that inference and prediction are separate tasks which should not be conflated in model selection (Shmueli 2010; Laubach *et al*. 2021; McElreath 2021). We echo Gelman and Rubin’s (1995) criticism of selecting “a model that is adequate for specific purposes without consideration of those purposes”. In the context of occupancy models, both explanation and prediction are important objectives, and conflating the two does justice to neither. Furthermore, our results emphasise the importance of considering not only the model’s purpose, but also the purpose of sub-models within the model; the purpose of the occupancy sub-model depends on whether we are interested in predicting the occupancy state or inferring its drivers, while the detection sub-model’s purpose is almost always prediction of the detection probability. Consequently, how occupancy covariates are chosen depends on the purpose of the model – information criteria are suitable if the purpose of the model is prediction, but are unlikely to be if the purpose is parameter inference – while detection covariates can generally be selected using information criteria. This advice also applies to other predictive model selection methods such as cross-validation; the choice of model made by AIC is asymptotically equivalent to that made by leave-one-out cross validation (Stone 1977).

### Using information to compare biological hypotheses in observational studies is risky

The importance of distinguishing between inference and prediction has wider implications for how information-theoretic model selection is applied in ecology. Proponents of the information-theoretic approach have argued that it can be used to compare multiple *a priori* specified models, each representing a different biological hypothesis, with the relative AIC scores indicating the strength of evidence for each hypothesis (Johnson & Omland 2004; Richards 2005; Burnham *et al*. 2011). However, using information criteria in this way conflates inference and prediction; information criteria select models which make better predictions, but these same models can contain spurious effect sizes which hold no biological meaning, while the effects of biologically important covariates are confounded. This is not only the case for occupancy models; the occupancy models we employed are just an extension of logistic regression (Clark & Altwegg 2019), and these points apply to other forms of linear model as well (Luque-Fernandez *et al*. 2019; McElreath 2021). The implication is that using information-theoretic model selection to compare biological hypotheses in observational studies carries substantial risks.

Guarding against the risks associated with using information criteria to compare biological hypotheses requires the use of subject expertise to make assumptions about the potential relationships between variables in the system, to determine if phenomena such as collider bias or post-treatment bias may be at play. These assumptions should be communicated clearly (*e*.*g*., as a DAG) so that they are open to critique and verification, and so that the implications of different assumptions can be compared (Pearl 2009). Additionally, we argue that if it is possible to make these assumptions, then methods such as the back-door criterion can be employed to choose a suitable set of covariates for the model to estimate the effect or effects of interest, circumventing the need for information criteria in these cases. Therefore, we argue that if the aim is to compare biological hypotheses to infer the drivers of species occurrence or to understand the consequences of management interventions, then causal inference is the more suitable approach.

### The information-theoretic approach and causal inference are complementary

While we argue that comparing biological hypotheses using the information-theoretic approach is risky, and that we prefer a causal inference-based approach for this purpose, we must emphasise that we are not arguing that the information-theoretic approach is flawed or useless for model selection. Information criteria select models from the “predictive point of view” (Akaike 1998), while causal inference is concerned with estimating the effects of covariates, so we see the two approaches as complementary. In the case of occupancy models the two approaches may be used side-by-side in a single analysis, where occupancy covariates are chosen based on causal assumptions embodied in a DAG, while the detection covariates are selected using the information-theoretic approach.

We also argue that causal inference and the information-theoretic approach are complementary because they share many philosophical underpinnings. Proponents of the information-theoretic approach emphasise the importance of subject expertise and *a priori* “hard thinking” to develop hypotheses which are then compared as models (Lukacs *et al*. 2007; Burnham *et al*. 2011). In causal inference, subject expertise and *a priori* thought are vital in making the causal assumptions which are embodied in the DAG (Pearl 1995; Greenland *et al*. 1999). Causal inference therefore provides a framework to support the “hard thinking” required in ecological modelling (Grace & Irvine 2020). Proponents of the information-theoretic approach also recognise that “a proper analysis must consider the science context and cannot successfully be based on ‘just the numbers’” (Burnham & Anderson 2004). Similarly, proponents of causal inference argue that conclusions cannot be drawn from the data alone, but require causal assumptions which come from the scientific context of the model (Pearl 2009; Pearl *et al*. 2016).

Chamberlin’s (1890) method of multiple working hypotheses is often emphasised in the information-theoretic approach (Burnham & Anderson 2004; Elliott & Brook 2007). We argue that causal inference is very compatible with Chamberlin’s method; constructing a causal model forces us to consider multiple explanations for a phenomenon, guarding against the threat of “parental affection for a favourite theory” which concerned Chamberlin. Furthermore, the causal approach, like the method of multiple working hypotheses, does not rely on the comparison of a model with a null model or hypothesis which is often false on *a priori* grounds and therefore provides little insight when it is rejected (Anderson *et al*. 2000; Burnham & Anderson 2004). Due to the relatively static nature of causal models, we argue they are especially suited to the case of multiple working hypotheses in parallel (Elliott & Brook 2007), in which causation operates through multiple factors simultaneously. Moreover, the tools of causal inference allow this parallel case to be extended to more complex situations with indirect effects, rather than constraining our thinking to simple additive terms and interactions.

Finally, proponents of the information-theoretic approach often emphasise George Box’s classic point that “all models are wrong, but some of them are useful” – they do not believe in true models (Burnham & Anderson 2004). We agree, and argue that causal inference does not involve making models that are true, but instead involves making models which are useful in the sense of inferring the drivers of patterns and predicting the effect of interventions. Causal inference provides the tools to express the assumptions underpinning these inferences in a transparent manner, and allows us to examine how robust our inferences are by using sensitivity analyses to explore the consequences of different causal assumptions and latent confounding variables.

### A caveat: model selection is more than selecting covariates

A caveat which must be borne in mind is that our article focuses on the choice of variables to condition on (i.e., include as covariates), which is a key aspect of model selection, but another vital part of model selection is selecting specific mathematical functions to relate these variables to one-another (Johnson & Omland, 2004). However, as the rules of causal inference are non-parametric in the sense that they do not assume any specific form in the functions relating variable (Pearl 1995; Greenland *et al*. 1999) our conclusions hold irrespective of what functional forms are chosen, and we consider any role of information criteria in selecting these functions to be beyond the scope of our article.

### Summary

We have demonstrated that when a form of collider bias known as M-bias is present in the occupancy process, occupancy models which are favoured by AIC and BIC produce inaccurate parameter estimates but accurate predictions. In contrast, M-bias in the detection process does not affect the accuracy of parameter estimates. The key conclusion supported by these results is that inference and prediction are separate tasks which should not be conflated during model selection. The correct choice of model selection procedure depends on the purpose for which the occupancy model will be used. Information-theoretic approaches are suitable for selecting occupancy covariates if the model is to be used for predicting the site-level occupancy probability. However, if the goal is instead to infer the effect of environmental covariates on occupancy, then the use of information criteria carries significant risks; we advocate for an approach based on causal inference in this situation. Our results support the selection of detection covariates using information-theoretic methods regardless of the model’s purpose, because detection probability is almost always a nuisance parameter. As the single-season occupancy models we used are in essence a form of logistic regression, our results have wider implications for the use of information-theoretic model selection in ecology. In particular, we argue that our results, alongside those of others (Luque-Fernandez *et al*. 2019; McElreath 2021), underscore the risks associated with using the information-theoretic approach to compare biological hypotheses in observational studies. Causal inference and the information-theoretic approach share similar philosophical underpinnings, and should be seen as complementary tools that accomplish different tasks.

## Acknowledgements

The authors would like to thank the members of the Conservation Ecology Group (Durham University) for their stimulating discussions which improved the content of this manuscript. P.S.S was supported by the Natural Environment Research Council (NERC) through the Iapetus2 Doctoral Training Partnership.

