## Supplementary figures for "Model Selection in Occupancy Models: Inference versus Prediction"

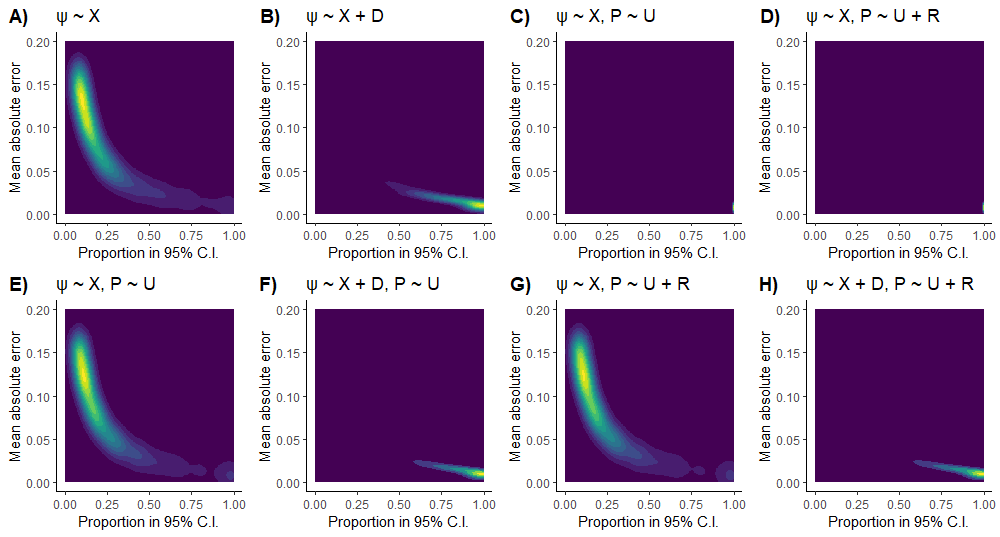

**Figure S1.** Kernel density estimate contours showing two measures of retrodictive accuracy when predicting site-level occupancy probability (ψ), for 1000 simulations. The x-axis shows the proportion of sites (out of 3000 total sites) for which the true occupancy probability was contained within the 95% confidence interval around the model’s prediction. The y-axis shows the mean absolute error of the predictions. Consequently, the bottom right of the plot indicates higher predictive accuracy, while the top left indicates lower predictive accuracy. The density of simulations within this area is shown by the coloured contours, with lighter colours indicating a higher density of simulations. Results are displayed for the following occupancy models: **A)** scenario 1, model 1; **B)** scenario 1, model 2; **C)** scenario 2, model 1; **D)** scenario 2, model 2; **E)** scenario 3, model 1; **F)** scenario 3, model 2; **G)** scenario 3, model 3; **H)** scenario 4, model 4. Each model’s covariates for ψ and the detection probability (P) are shown above their respective plot. For explanation of the scenarios, see Table 1 and main text.

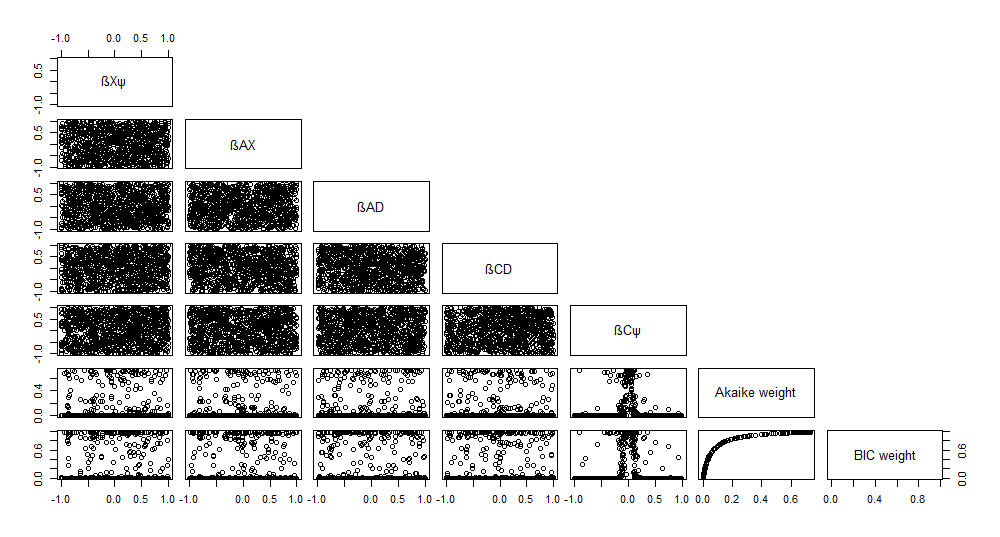

**Figure S2.** Pairs plot showing the relationship between the effect sizes (β values), Akaike weight, and BIC weight for model 1 in scenario 1 (see Table 1 for model definition).

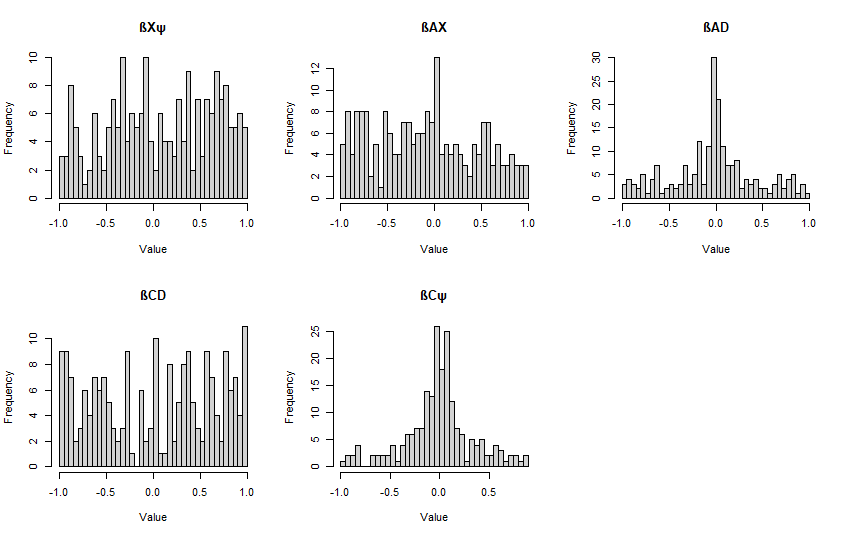

**Figure S3.** Histograms showing distribution of true parameter values for the simulations in scenario 1, model 2 (see Table 1 for model definition) for which the true effect of X on ψ (β_Xψ_) was within the 95% confidence interval around the estimate.

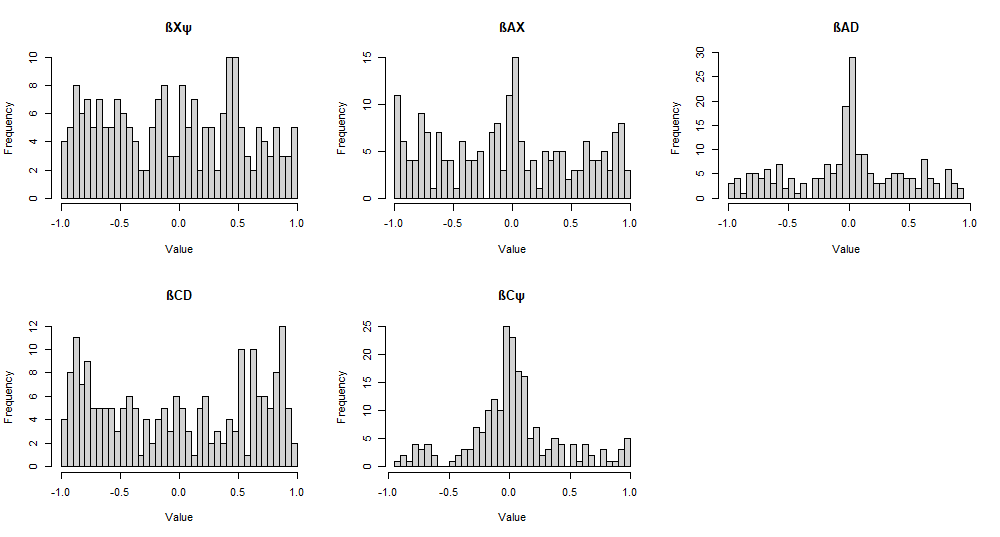

**Figure S4.** Histograms showing distribution of true parameter values for the simulations in scenario 3, model 2 (see Table 1 for model definition) for which the true effect of X on ψ (β_Xψ_) was within the 95% confidence interval around the estimate.

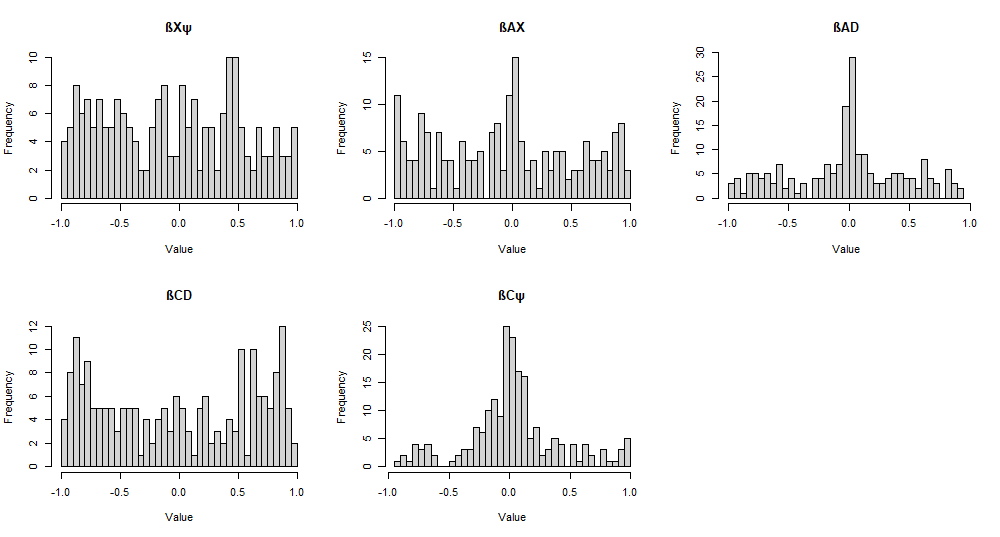

**Figure S5.** Histograms showing distribution of true parameter values for the simulations in scenario 3, model 4 (see Table 1 for model definition) for which the true effect of X on ψ (β_Xψ_) was within the 95% confidence interval around the estimate.

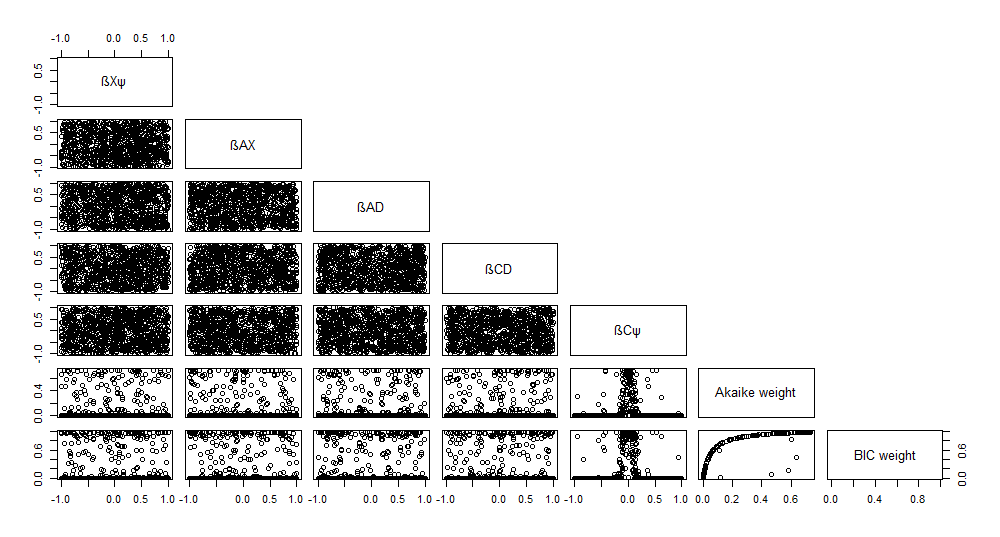

**Figure S6.** Pairs plot showing the relationship between the effect sizes (β values), Akaike weight, and BIC weight for model 3 in scenario 3 (see Table 1 for model definition).
